# Animal product-free formation and cultivation of 3D primary hepatocyte spheroids

**DOI:** 10.1101/2025.04.10.648088

**Authors:** Evgeniya Mickols, Lazaros Primpas, Stina Oredsson, Maria Karlgren

## Abstract

3D cultures of primary human hepatocytes (3D PHH) are successfully used to reduce and replace the use of animal experiments in biomedical research. Yet, the initial formation of 3D PHH is highly dependent on the supplementation with fetal bovine serum (FBS). However, the molecular composition of FBS and its effects on cultured cells are poorly understood. Moreover, FBS is prone to batch-to-batch variation, immunogenic risk and lack of adherence to the replacement, refinement, and reduction (3Rs) of animal experiments. Here, we demonstrate that FBS can be fully replaced by animal-free substitutes, thus facilitating fully chemically defined and animal serum-free 3D PHH cultures. Specifically, we combined a previously developed animal-free substitute cocktail (Rafnsdóttir *et al*., 2023) with a normoglycemic (5.5 mM glucose and 0.58 ng/mL insulin) chemically defined culture medium (Handin *et al*., 2023). Morphological and viability evaluations, along with global proteomics data, demonstrated that serum-free cultured 3D PHH have equal or superior viability and functional performance of cytochrome P450s, rendering this medium useful for long-term studies and *in vitro* ADMET applications. This study marks a significant advancement in the development of animal serum-free culture conditions for primary human cell cultures, paving the way for more reliable and ethical *in vitro* studies.

**Significance statement:** Most *in vitro* cell models rely on fetal bovine serum (FBS). However, the use of FBS leads to inconsistent experimental results and raises serious ethical concerns. In our study, we develop and evaluate chemically defined animal product-free cell culture medium with physiologically relevant levels of key hormones and nutrients for liver spheroid cultures. This study marks a significant advancement in the development of animal serum-free culture conditions for primary human cell cultures used in drug disposition studies.

## Introduction

*In vitro* primary human cells cultures are successfully applied to reduce and replace the use of animal experiments in biomedical research (Wang et al., 2024). Of note, relevant *in vitro* models even offer the potential to accelerate drug development, and are increasingly incorporated into regulatory preclinical safety assessments. (Turner et al., 2023). Although these *in vitro* models greatly contribute to the replacement, refinement, and reduction (3Rs) of animal experiments, culturing conditions for most *in vitro* models include the animal-derived component fetal bovine serum (FBS).

The use of FBS is a routine cell culture practice. It has been used for the maintenance of cell cultures for nearly 70 years, and it remains the most common cell culture medium supplement (Puck et al., 1958). FBS is added to the culture medium to help the cells circumvent stress induced by the *in vitro* culture setting. However, the molecular composition of FBS is prone to batch-to-batch variation and is poorly understood at the molecular level (Gstraunthaler et al., 2013; Baker, 2016; Liu et al., 2023). In addition, the use of FBS does not adhere to the 3Rs, as FBS production is associated with severe animal welfare issues. (Valk et al., 2018).

Although the problems associated with FBS are well-known, limited research has been dedicated to finding serum-free alternatives for *in vitro* models. Nonetheless, in recent studies focusing on serum replacement, the authors demonstrate that FBS-free media could be successfully developed and used to support *in vitro* cultures of various cell lines (Stephenson et al., 2012; Edwards et al., 2018; Fiol et al., 2023; Perez-Diaz et al., 2023; Rafnsdóttir et al., 2023; Dai et al., 2024; Katayama et al., 2024; Weber et al., 2024). To substitute FBS one would need to supplement cell culture medium with either human blood derivatives or a cocktail of human-derived or recombinant attachment, growth and carrier proteins as well as minerals, hormones and vitamins. Such cell culture media are available as proprietary solutions from life science supplier, however the content of these media is undisclosed, which makes customization of the media or interpretation of the research results tricky. Alternatively, Rafnsdóttir *et al*. recently published the composition of a fully defined animal-product free and universal medium that could be used to effectively substitute FBS in long-term cultures in adherence to the 3Rs (Rafnsdóttir et al., 2023). As opposed to the proprietary cell culture media, the approach provided by Rafnsdóttir *et al*. is fully disclosed and well-described, thus opening an opportunity for customization of serum-free medium (SFM) and adaptation of the basal components to the cell culture of choice. In addition, the paper by Rafnsdóttir *et al*. has recently been supplemented with a detailed methodological paper covering all aspects of the animal serum-substitute preparations (Weber et al., 2024).

In this paper, we focus on developing and evaluating FBS-free 3D primary human hepatocyte (3D PHH) cultures for drug disposition studies. 3D PHH have recently emerged as a gold standard model for prediction of liver toxicity and drug clearance (Bell et al., 2016, 2018; Vorrink et al., 2018; Handin et al., 2021; Kanebratt et al., 2021; Preiss et al., 2022; Mickols et al., 2024). Notably, from a 3Rs perspective, 3D PHH are nearly an ideal *in vitro* model — it is sourced from surplus human tissue, cultured in high-throughput microwell format in a chemically defined medium with minimal supplementation. Yet, the initial formation of 3D PHH has so far been highly dependent on the supplementation of the medium with FBS. This decreases the 3Rs compliance of 3D PHH, and possibly, tampers with the desired ADMET-related properties, due to FBS batch-to-batch variability. Thus, FBS-free 3D PHH formation is desirable. Importantly, we previously demonstrated that 3D PHH can be cultured in chemically defined physiologically relevant culture medium with fasting levels of glucose and insulin for at least two to three weeks (Handin et al., 2021; Mickols et al., 2024). Importantly, this condition supports the hepatic phenotype and the expression of absorption, distribution, metabolism, excretion, and toxicity (ADMET) related proteins and their functions.

Here, we transition the 3D PHH system to a fully animal serum-free physiologically relevant model and comprehensively benchmark its performance and utility for preclinical pharmacological applications. Specifically, we compared 3D PHH spheroid formation, viability, cytochrome P450 (CYP) activity, and global proteome of spheroids formed in an FBS-supplemented medium with those formed in the fully defined serum-free condition. We demonstrate that chemically defined animal serum-free culture could replace FBS and improve 3D PHH culture stability and CYP metabolic function.

## Materials and methods

### Isolation of primary human hepatocytes

Human hepatocytes were isolated from histologically normal surplus liver tissue obtained from cancer patients undergoing liver resections at the Department of Surgery at Uppsala University Hospital. Donors signed informed consent in agreement with the approval from the Uppsala Regional Ethical Review Board (Ethical Approval no. 2009/028, amended 2018/1108). A previously described two-step perfusion procedure for hepatocyte isolation was performed (Lecluyse and Alexandre, 2010). Briefly, the liver tissue was rinsed from excessive blood with Hypothermosol FRS (Biolife Solutions, Bothell, WA) and perfused with collagenase and protease buffers for tissue digestion. Isolated hepatocytes were further centrifuged with isotonic Percoll for debris removal and cryopreserved until further use as previously described (Ölander et al., 2019, 2020, 2021).

### 3D Primary human hepatocyte spheroid culture

Five PHH donors were used in this study (Supplementary Table 1). Cryopreserved hepatocytes were gently thawed and transferred to isotonic 27 % Percoll (GE Healthcare, Chicago, Illinois) in Williams E normoglycemic medium (PAN-Biotech GmbH, Aidenbach, Germany) without serum (from here and further referred to as WEng; full medium composition is provided in Supplementary Table 2) and centrifuged at 100 x *g* for 10 minutes (Handin et al., 2021; Mickols et al., 2024). After the centrifugation, the supernatant with cell debris and dead cells in Percoll was discarded. The hepatocytes were resuspended in a small volume of warm WEng medium and cell viability and count were measured with acridine orange−propidium iodide staining using a Cellometer Vision CBA image cytometer (Nexcelom Bioscience, Lawrence, USA). Then, hepatocytes were resuspended either in WEng medium supplemented with 10 % FBS or with the serum substitute, containing e.g. recombinant hormones, proteins, growth factors etc., developed by Rafnsdóttir *et al* (Rafnsdóttir et al., 2023a). Full composition of the serum substitute is provided in Supplementary Table 2. The PHH suspensions were seeded at 2,000 cells/well in 100 µl of cell culture medium in ultra-low attachment 96-well plates Corning 7007 (Corning, Kaiserslautern, Germany), sedimented by 100 x *g* centrifugation for 5 minutes, and incubated at 37°C in a humidified incubator with 5 % CO_2_ in air (Figure 1a). The first medium change was typically performed after the initial spheroid formation (5-7 days after the seeding of PHHs), starting with a 50 % medium change to WEng without any additional serum/serum-substitute supplement. Spheroids were maintained in unsupplemented WEng for up to 3 weeks with a medium change every 48 to 72 h. All media and cell-culture supplements were purchased from VWR (Radnor, Pennsylvania), Thermo Fisher Scientific (Waltham, Massachusetts), or Sigma-Aldrich (Saint Louis, Missouri) unless otherwise stated. The serum substitute cocktail developed by Rafnsdóttir *et al*. was prepared according to Weber *et al*. in Prof. Oredsson’s laboratory, Lund University, and the composition of the supplement is provided in Supplementary Table 2 (Rafnsdóttir et al., 2023a; Weber et al., 2024).

**Figure 1.**
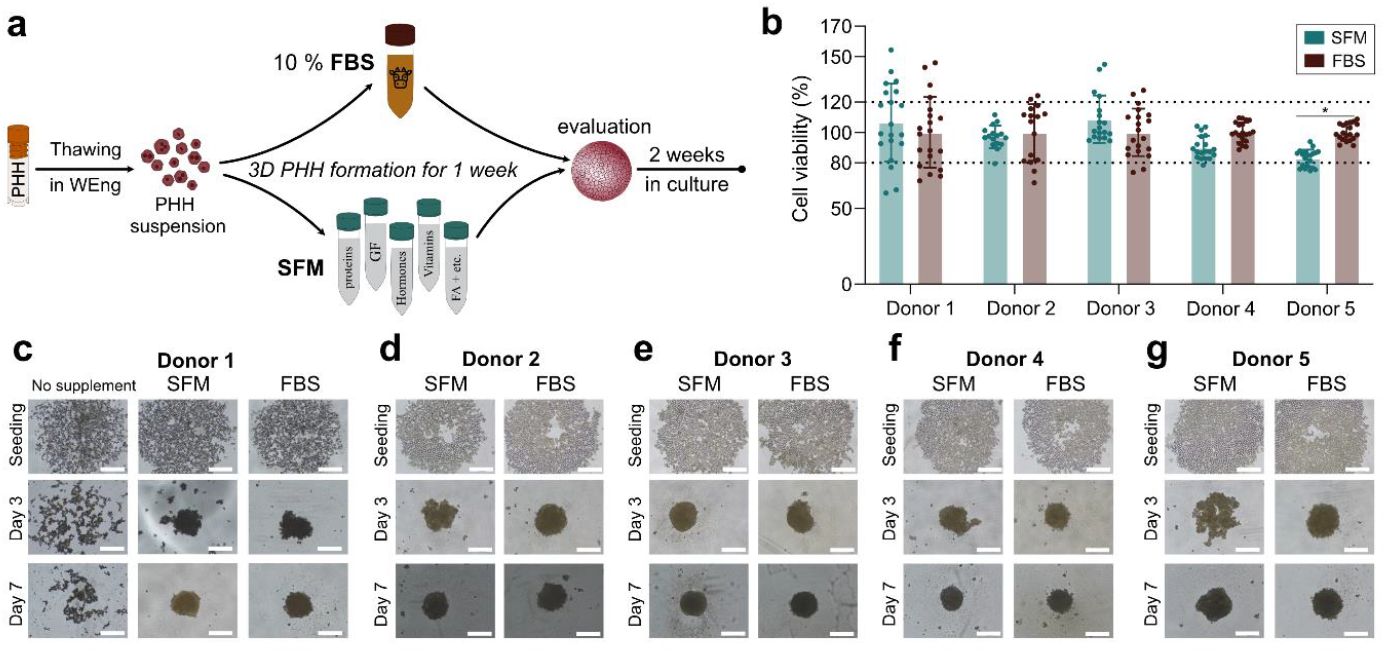
3D primary human hepatocyte spheroid (3D PHH) formation in serum-free medium (SFM) or conventional fetal bovine serum (FBS)-supplemented medium. **a**. Schematic overview of the 3D PHH seeding and culture maintenance procedures. PHH were thawed in William’s E normoglycemic medium (WEng) and seeded with 10 % FBS or serum substitute according to Rafnsdottir *et al* (Rafnsdóttir et al., 2023). After full spheroid formation (one week) culture medium was exchanged to unsupplemented WEng. **b**. Viability (ATP content) of 3D PHH from five different donors in SFM or with FBS supplementation one week after seeding. Data are presented as mean ± SD for viability measurements normalized to conventional conditions with FBS supplementation (n=18-20). The dotted lines indicate viability ratios of 80 to 120 %. *, p < 0.005; in two-way ANOVA with Šídák’s multiple comparisons test. **c**,**d**,**e**,**f**,**g**. 3D PHH formation in five donors. Bright-field microscopy images were taken at seeding, after the third day in culture and after one week in culture. Scale bar = 250 µm.

### PHH viability assay

PHH viability/health state was assessed using the CellTiter-Glo® 3D assay (Promega, Madison, Wisconsin) according to the manufacturer’s instructions. Briefly, the 3D PHH plates and CellTiter-Glo® 3D reagent were equilibrated to room temperature for approximately 30 minutes, and the CellTiter-Glo® 3D reagent was added to the well in a volume equal to the cell culture medium. The plates were incubated on an PST-60HL-4 Plate shaker-thermostat (Riga, Latvia) for approximately 30 minutes to facilitate the disintegration of the spheroids. The luminescence was then measured in a TECAN Spark plate reader (Zürich, Switzerland) for a minimum number of sixteen technical replicates/spheroids.

### Cytochrome P450 activity

Single 3D PHH were incubated for 8 h at 37°C in WEng medium with a cocktail of CYP substrates: 10 μM midazolam (CYP3A4), 10 μM diclofenac (CYP2C9), 10 μM bufuralol (CYP2D9) and 10 μM bupropion (CYP2B6). After 8 h, the reactions were stopped by mixing and snap-freezing the samples. For analysis, samples were thawed and diluted with acetonitrile (sample:acetonitrile - 60:40) supplemented with 10 nM warfarin as an internal standard. Samples were centrifuged twice for 10 min at 4°C with 3500 × *g* to sediment cell pellets and proteins, and concentrations of the respective CYP metabolites (1-hydroxymidazolam, 4-hudroxydiclofenac, 1-hudroxybufuralol and hydroxybupropion) in supernatants were analyzed on a Waters Acquity UPLC coupled to Waters Xevo TQ MS with electrospray ionization (Liquid chromatography conditions and MS/MS transitions are provided in Supplementary Tables 3 and 4 respectively). Compounds were separated with a 2 min gradient elution of acetonitrile and 0.1 % formic acid (flowrate 0.5 mL/min) on a Waters BEH C18 column, 2.1 × 50 mm (1.7 μm) at 60°C. Peaks were quantified using MassLynx Software V4.2 with TargetLynx module.

### Global proteomics analysis

#### Sample preparation procedure

3D PHH (at minimum 100 spheroids per condition and three biological replicates) were collected in low protein binding microcentrifuge tubes (Eppendorf, Fisher Scientific, Gothenburg), washed twice with ice-cold PBS, and snap-frozen in liquid nitrogen. Then samples were rapidly defrosted and proteins were prepared using the iST sample preparation kit (PreOmics, München, Germany) according to the instruction manual with the exception of the reconstitution buffer that was exchanged to 0.1 % formic acid in LC-MS/MS grade water, transferred to glass LC-MS vials and dried in a Speedvac at 40°C. Final peptide concentrations were determined by fluorimetric quantification with the Protein Broad Range Assay on the Qubit 4 Fluorometer (Thermo Fisher Scientific, Waltham, MA).

#### LC-MS/MS analysis of the peptides

Peptides were separated on an EASY-spray C18-column (50 cm, 75 μm inner diameter), using an acetonitrile/water gradient (0.1 % formic acid) at 300 nL/min. Eluted peptides were analyzed using the TopN method (full MS followed by ddMS2 scans) on an Orbitrap Q Exactive HF mass spectrometer (Thermo Fisher Scientific), operating in a data-dependent mode with survey scans at a resolution of 120,000, AGC target of 3 × 10^6^ and maximum injection time of 120 ms. The top 15 most abundant isotope patterns were selected from the survey scan with an isolation window of 1.7 m/z and fragmented with nCE at 26. The MS/MS analysis was performed with a resolution of 30,000, AGC target of 1 × 10^5^ and a maximum injection time of 50 ms. Blanks were injected between every sample to ensure minimal transfer of peptides between biological replicates and different conditions. The mass spectrometry proteomics data have been deposited to the ProteomeXchange Consortium via the PRIDE partner repository with the dataset identifier PXD056428 (*reviewer login details username are available upon request*). (Perez-Riverol et al., 2021, 2019)

#### Data analysis

The raw MS datafiles were processed using MaxQuant version 2.5.2.0. (Cox and Mann, 2008) Proteins were identified by searching MS and MS/MS data of peptides against a reference human proteome database retrieved from the UniProtKB/Swiss-Prot curated database on 2024-05-11 and contaminants FASTA provided by MaxQuant (The UniProt consortium, 2024.). Description of the parameters used for peptide identification by MaxQuant can be found in the mqpar.xml uploaded to PRIDE. Briefly, carbamidomethylation was set as fixed modification, while oxidation and acetylation as variable modifications; match between runs algorithm was used; decoy sequences were created by reversing the target sequences and peptide-spectrum matches, peptides and proteins were validated at a 1 % false discovery rate estimated using the decoy hit distribution. Quality control of the MaxQuant search was performed using an R-based quality control pipeline called Proteomics Quality Control (PTXQC) version v1.0.16 (Bielow et al., 2016).

Subsequent data clean-up was performed using an in-house developed proteomics data pipeline in development, R version 4.3.0. Concisely, the ProteinGroups data table was cleaned up using Tidyverse package, and raw intensities were normalized using variance stabilization normalization (Vsn) using vsn package (Bioconductor 3.17 release) (Välikangas et al., 2018; Huber et al., 2002). Protein abundances (fmol/μg total protein) were calculated with the total protein approach (TPA) (Wiśniewski et al., 2014). Data overview analysis was performed in Perseus version 2.0.10.0 (Tyanova et al., 2016). Differential expression (DE) analysis was implemented via Amica web-platform version 3.0.1 (Didusch et al., 2022). Gene ontology enrichment analysis was performed using the GOrilla tool (Eden et al., 2007, 2009). Normalised intensities table, protein abundances suing TPA and results of DE are summarized in Supplementary data.

#### Quantification and statistical analysis

Unless otherwise stated, statistical analysis and plot generation were carried out using GraphPad Prism version 10 (GraphPad Software, San Diego, CA). All results are presented as mean values ± standard deviations of at least three biological replicates if not otherwise specified.

## Results

### Spheroid formation in animal serum-free medium

To be used in an experimental setting, PHH need to successfully assemble into spheroids. Since extracellular matrix and transmembrane cell adhesion proteins are typically lost due to the digestion associated with the PHH isolation procedure (Handin et al., 2021), it is nearly impossible for 3D PHH to self-assemble without additional structural proteins with preserved phenotype in a reasonable experimental time (Supplementary Figure 1). To circumvent this limitation and combat additional cell stress associated with cryopreservation, PHH are typically thawed and seeded in a medium supplemented with FBS (Bell et al., 2016; Handin et al., 2021; Mickols et al., 2024).

Here we used a humanized serum-substitute cocktail developed by Rafnsdóttir *et al*. to substitute FBS during the spheroid formation time (Figure 1a). Intriguingly, PHH efficiently assembled into well-defined viable 3D spheroids in the SFM by the seventh day in culture (Figures 1b-g). We observed this successful 3D PHH formation in SFM across five different donors, yet the speed of this process and morphology of spheroids in both SFM and FBS varied (Figures 1c-g). Thus, we asked five 3D PHH experts, that are not involved in the current study, to blindly assess the morphology of the 3D PHHs formed in SFM or FBS after one week of culture time. (Supplementary Tables 5 and 6). As expected, 3D PHH from different donors were scored differently (average score 2.9 to 4.1). However, 3D PHH formed in SFM obtained consistently higher or similar scores compared to the scores for 3D PHH formed in FBS, further confirming the successful microtissue formation in FBS-free conditions.

Additionally, to assess the formed 3D PHH from five biological donors, we performed ATP content measurement as an estimation of viability one week after the seeding (Figure 1b). For all five PHH donors, the viability of the 3D PHH formed in SFM stayed within control levels *i*.*e*. within 80 to 120 % change normalized to the conventional FBS setting. For donor 5, a statistically significant reduction in viability was seen for the 3D PHH formed in SFM (Figure 1b). Nonetheless, the viability was still within the acceptance limit (82 % of the control values).

### Long-term cultures

As 3D PHH are often used for long-term cultures, we randomly selected one of the five PHH donors (Donor 1). PHH from that donor were seeded again in SFM or in FBS-supplemented medium, and these 3D PHH cultures were used to ensure that animal serum-free conditions were applicable for long-term studies.

### Spheroid morphology

For long-term studies, we performed further evaluations with 3D PHH from donor 1 through three weeks in culture – a typical timeline for experiments involving this *in vitro* model (Figure 2a) (Kozyra et al., 2018; Vorrink et al., 2018; Handin et al., 2021; Koutsilieri et al., 2024; Mickols et al., 2024). We observed that the successfully formed spheroids preserved their morphology and integrity for the whole duration of the experiment (3 weeks), independently of whether the 3D PHH were formed with the support of FBS or in SFM.

**Figure 2.**
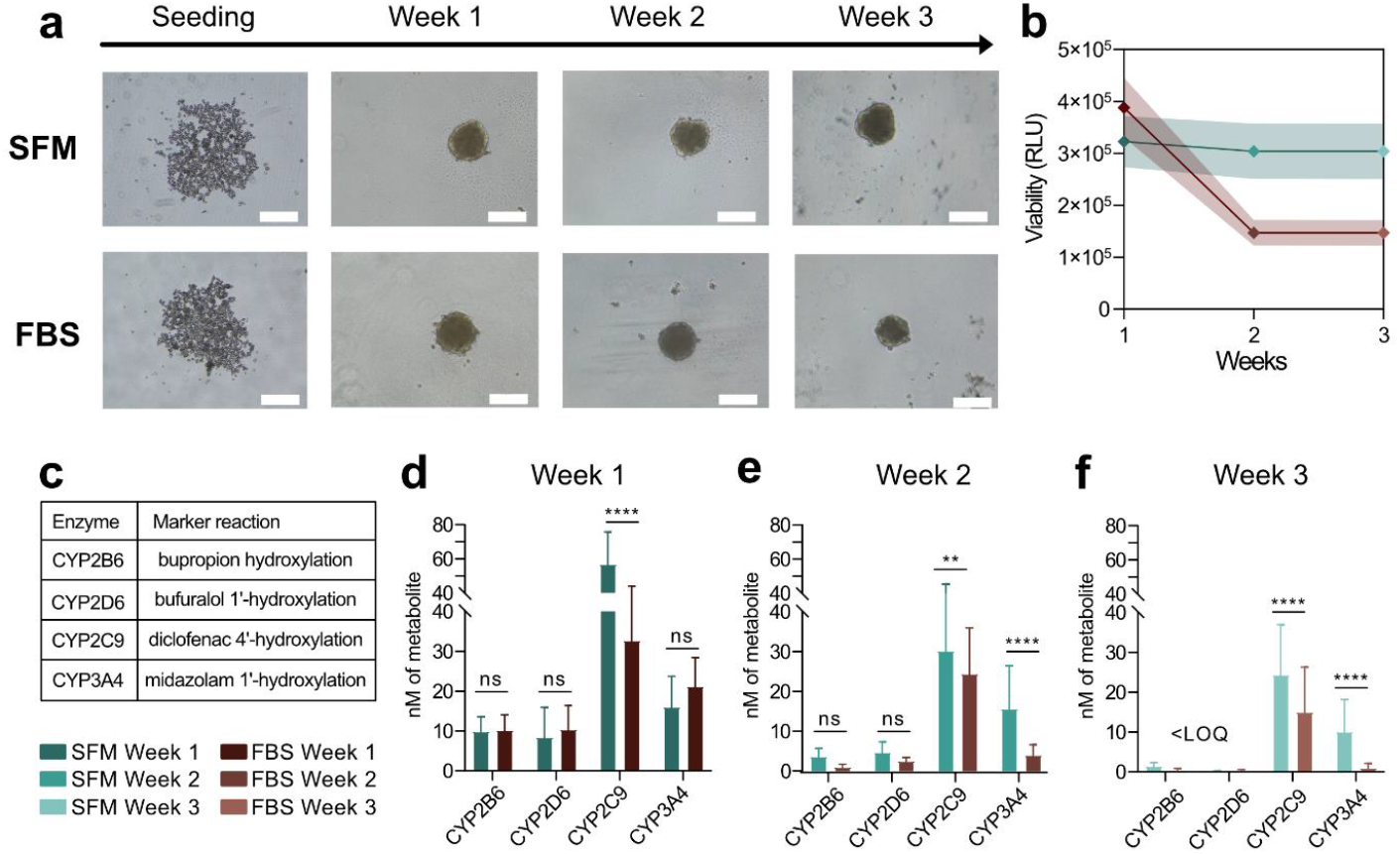
3D primary human hepatocyte spheroid (3D PHH) formation and function for three weeks. **a**. 3D PHH spheroid formation in serum-free medium (SFM) or fetal bovine serum (FBS)-supplemented medium and corresponding morphology for three weeks in culture. Scale bar = 250 µm. **b**. Mean viability (ATP content) for three weeks in culture in SFM (green) or FBS-supplemented medium (brown), the shaded area represents standard deviation (n=18-24). **c**. Marker reactions used for assessing cytochrome P450 (CYP) function. **d**,**e**,**f**. Marker metabolite formation in nM in spheroids cultured for one, two, and three weeks. Data are presented as mean ± standard deviation (n=19-39). **, p < 0.01; ****, p < 0.0001; ns, not significant using two-way ANOVA with Šídák’s multiple comparisons test. <LOQ signifies metabolites levels below the limit of quantification. All experiments were performed in Donor 1 (see Figure 1c).

### Viability of long-term cultures

Likewise, we measured viability in 3D PHH from donor 1 over the culture period of three weeks (Figure 2b). Interestingly, whilst ATP measurements were rather comparable between FBS- and SFM-formed 3D PHH during the first week of culture, a notable decrease (2-fold) in ATP quantities was seen for spheroids formed in FBS by the second week in culture. Meanwhile, ATP values remained stable in 3D PHH formed in SFM for the whole duration of culturing.

### Cytochrome P450 activity

3D PHH are increasingly used in drug disposition and metabolism studies (Vorrink et al., 2018; Hendriks et al., 2020; Kanebratt et al., 2021; Preiss et al., 2022; Järvinen et al., 2023). To validate the applicability of 3D PHH formed in SFM for such studies, we measured the activities of common drug metabolizing CYP enzymes after one, two and three weeks post-seeding (Figure 2c-f). We did not observe any difference in the activities of CYP2B6 and CYP2D6 enzymes between 3D PHH formed with FBS-supplemented medium or in SFM. However, metabolite formation by CYP2C9 was significantly higher in 3D PHH formed in SFM. Further, we observed that CYP3A4 activity was not significantly different at week 1 for the two different conditions (Figure 2d), whereas from the second week in culture and onwards spheroids formed in SFM exhibited significantly higher activity of CYP3A4 (Figures 2e,f).

### Global protein expression

Next, we investigated whether the use of SFM during the 3D PHH formation affected the proteomes of 3D PHH. In total, 4506 proteins were identified, and to obtain a global view of the dataset we applied principal component analysis (PCA). PCA indicated a clear, and expected, separation between proteomes of uncultured PHH in suspension and in 3D spheroid format with the first component explaining 24.8 % variance (Figure 3a) (cf. Handin et al., 2021). Interestingly, along the second component (13 % variance), the data separated mostly based on the culture time, and the difference between proteomes of 3D PHH formed in the two different conditions was not a driving factor of the separation (Figure 3a). The PCA data indicated that the shift from the conventional FBS-based 3D PHH protocol to the SFM protocol did not have a major impact on the proteomics fingerprint of 3D PHH *in vitro* cultures. However, we still observed that 3D PHH formed in these two conditions formed sub-clusters on the PCA plot.

**Figure 3.**
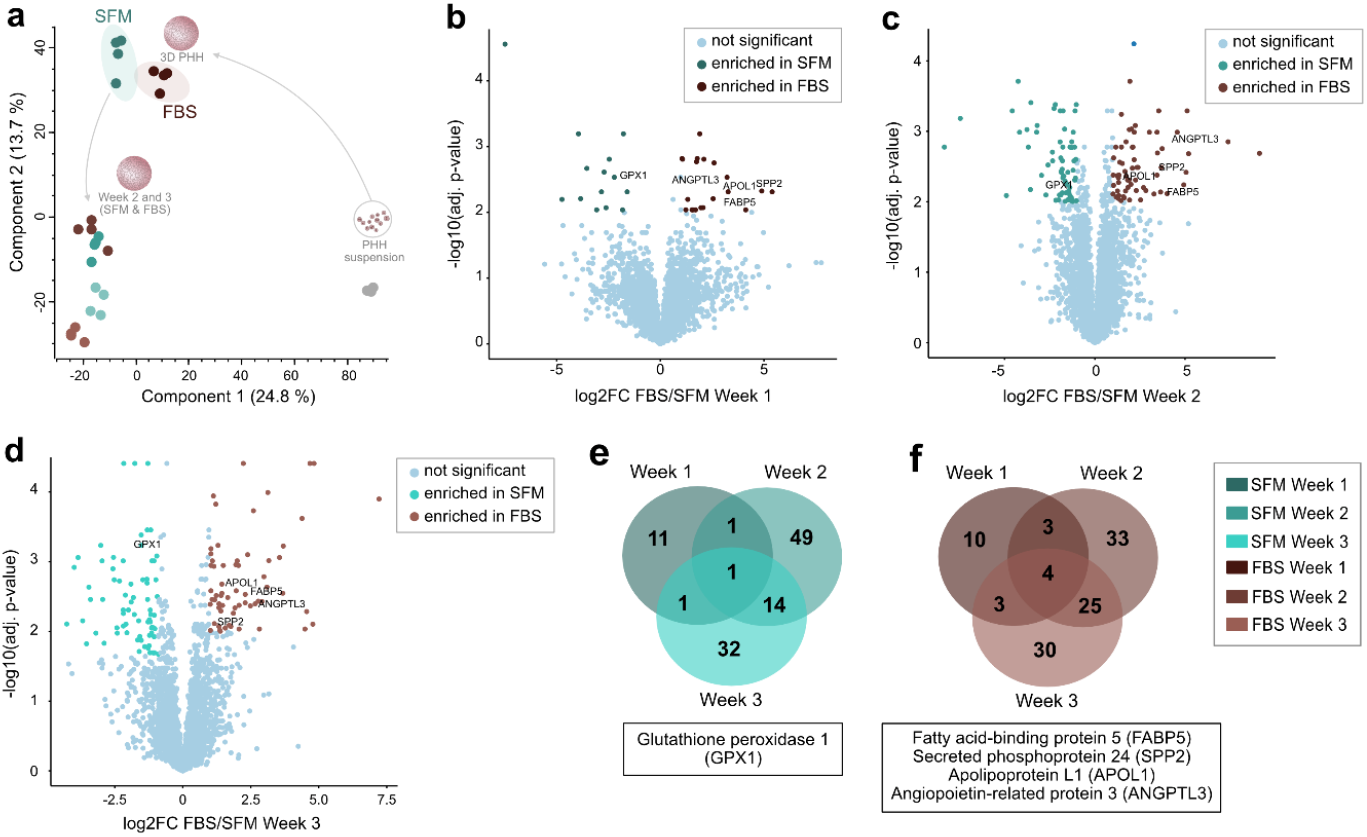
Proteomes of primary human hepatocyte spheroids (3D PHH) formed in serum-free medium (SFM) or fetal bovine serum (FBS)-supplemented medium. **a**. Principal component analysis of the global proteomes for the three culture weeks based on all identified proteins (4 biological replicates per condition, 4506 identified proteins). The number in parentheses is the percentage of variability explained by each component. **b**,**c**,**d**. Differential expression analysis (DeqMS) for 3D PHH cultures at week 1, 2 and 3 respectively. Log2 fold change-threshold = 1; significance threshold for multiple-hypothesis adjusted p-value = 0.01. **e**,**f**. Venn diagrams of the enriched differentially expressed proteins between 3D PHH formed with SFM or with FBS for one, two and three weeks. Consistently enriched differentially expressed proteins for SFM and FBS (panels e and f respectively) are highlighted in the boxes underneath the Venn diagrams and in panels b, c and d. (Proteomes correspond to Donor 1 cultures shown in Figure 2a.)

To further investigate possible differences between the conditions, we performed differential expression analysis (Figure 3b-d, Supplementary data). During the first week of 3D PHH cultures, we detected 34 differentially expressed proteins (DeqMS, Log2 fold change-threshold = 1, significance threshold for multiple-hypothesis adjusted p-value = 0.01). Unexpectedly, we detected more differentially expressed proteins by the second week post-seeding —130 proteins at the same statistical cut-off (Figure 3c, Supplementary data DE W1). We found no enriched pathways in 3D PHH formed in SFM, but observed multiple biological processes to be significantly altered in 3D PHH formed in FBS-supplemented medium (Supplementary data DE W2), e.g. negative regulation of catalytic activity (q=9.42E-5, e= 4.89) and negative regulation of molecular function (q=1.62E-3, e= 3.79). Likewise, after three weeks in culture, we found 110 protein to be differentially expressed (Figure 3d), and observed significant changes only in the 3D PHH formed in FBS-supplemented medium, that again exhibited negative changes in catalytic activity and other similar metabolic changes (Supplementary data DE W3). Nonetheless, almost no differentially expressed proteins across all conditions were connected with meaningful alterations in specific hepatic functions. Furthermore, we observed only a minor portion of the proteomes to be differentially expressed (0.8, 2.9 and 2.4 percent for weeks one, two and three, respectively; Figures 3b-d). Also, out of all differentially expressed proteins, only glutathione peroxidase 1 (GPX1) was consistently upregulated in spheroids formed in SFM during the whole duration of culture time (Figure 3e), whereas in 3D PHH formed in FBS-substituted medium, fatty acid-binding protein 5 (FABP5), secreted phosphoprotein 24 (SPP24), apolipoprotein L1 (APOL1) and angiopoietin-like 3 (ANGPTL3) were consistently upregulated (Figure 3f) when compared to SFM.

### Expression of clinically relevant ADMET proteins

To further support the applicability of 3D PHH formed in SFM in drug development studies, we analyzed the expression levels of the clinically relevant ADMET proteins mentioned in the recently released IHC M12 guidelines on drug interaction studies (Figure 4) (ICH M12 Guideline on drug interaction studies, 2024). Overall, the expression of ADMET proteins across all conditions was stable. We found no statistical difference in CYP expression levels, except for lower expression of CYP3A5 in spheroids formed in SFM during the first week of *in vitro* culture (Figure 4a). Likewise, we did not detect any significant expression differences for the P-glycoprotein (P-gp/MDR1), organic anion transporting polypeptide (OATP) 1B1 and 1B3 transporters (Figure 4b).

**Figure 4.**
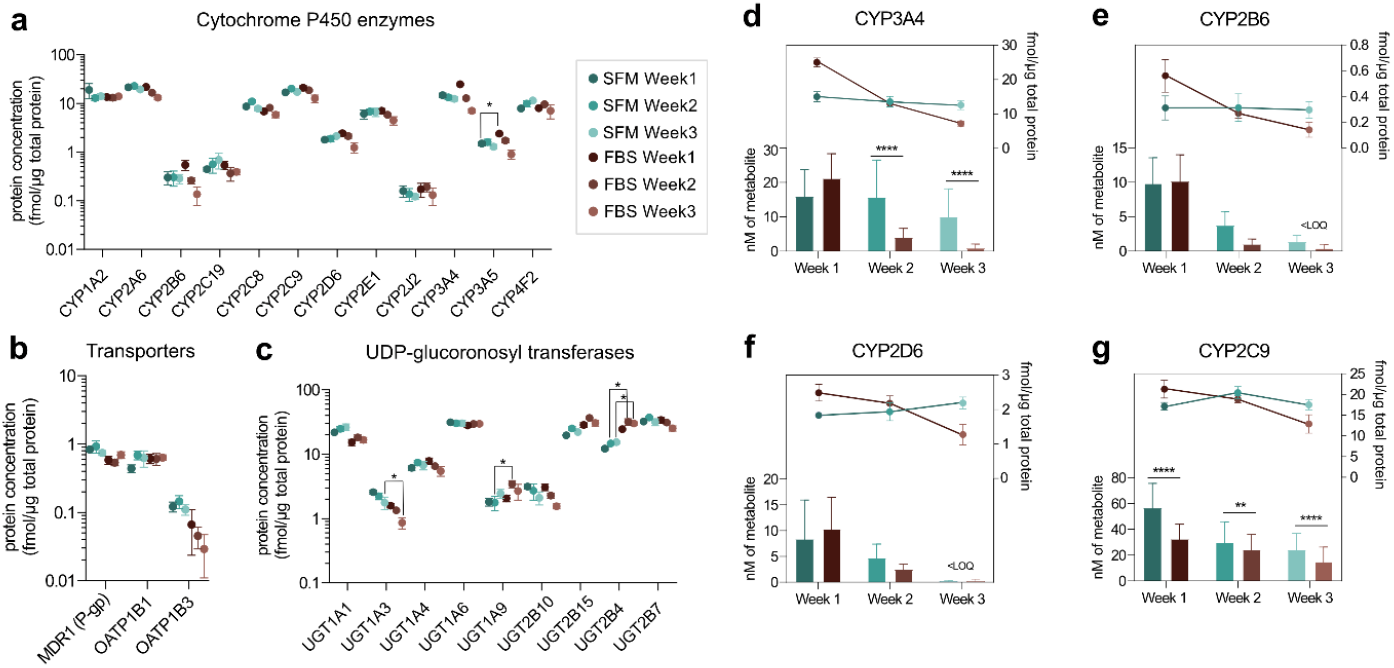
Global proteomics profiling of the ADMET-related proteins highlighted in ICH M12 guidelines. **a**. Expression levels of quantified cytochrome P450 (CYP) drug-metabolizing enzymes. **b**. Expression levels of quantified uridine diphosphate (UDP)-glucuronosyl transferases (UGTs). **c**. Expression levels of quantified drug transporters. Datapoints in all graphs represent mean expression values of four biological replicates with standard deviation. Stars signify differentially expressed proteins in differential expression analysis (DeqMS). **d-g**. Side-by-side comparison of expression and activity of CYP3A4, CYP2B6, CYP2D6 and CYP2C9, respectively.

Interestingly, among the ADMET proteins, we found most statistically significant differences in uridine 5’-diphosphate-glucuronosyltransferases (UGTs) expression with UGT1A3 being upregulated in 3D PHH formed in SFM during the third week of culture, whilst UGT1A9 and UGT2B4 were downregulated in the same condition during the second or second and third week of culture, respectively (Figure 4c).

Lastly, we compared the expression levels and the obtained activity of CYPs (Figure 4d-g). Similarly, to what has been demonstrated by others, we observed that in conventional FBS-based conditions, CYP activity and expression decreased in 3D PHH along the culture time (Figure 2d-f, Figure 4d-g) (cf. Handin et al., 2021; cf. Kanebratt et al., 2021). However, we noted that for this donor 3D PHH formed in SFM preserved a stable expression of CYPs over three weeks of culturing. In addition, we observed CYP activity to be notably more stable in 3D PHH formed in SFM, in particular for CYP2C9 and CYP3A4 (Figures 2d-f, Figures 4 d and g).

## Discussion

While earlier studies have explored the possibility of ADMET models being introduced at least temporarily to serum-free conditions, (Ranaldi et al., 2003; Bell et al., 2016; Rodrigues et al., 2020; Gheytanchi et al., 2021; Handin et al., 2021; Kanebratt et al., 2021; Ross et al., 2021; Mickols et al., 2024) fully chemically defined 3D PHH cultures, from the PHH seeding until the culture termination, have not been evaluated. Here, we utilized the serum-free cocktail developed by Rafnsdóttir *et al*., and followed up on the series of papers from our laboratory that demonstrate excellent PHH performance in a physiologically relevant culture medium supplemented with fasting levels of insulin and glucose (Handin et al., 2021; Rafnsdóttir et al., 2023; Mickols et al., 2024).

In all standard 3D PHH culture protocols, FBS is used at seeding. (Bell et al., 2016; Handin et al., 2021; Mickols et al., 2024; Oliva-Vilarnau et al., 2020; Oliva-Vilarnau et al., 2023; Xing et al., 2024) This as adhesion proteins and supplements present in FBS appear to be necessary for spheroid formation. Thus, an addition of FBS (or serum substitute) seems crucial for PHH self-assembly. Also, here we show that PHH seeded completely without any supplements do not easily or just never form spheroids (Figure 1c, Supplementary figure 1). This could be explained by the natural synthesis of structural proteins by the liver, which the hepatocytes are responsible for in the body. In contrast, PHH seeded in a medium supplemented with the serum-free cocktail suggested by Rafnsdóttir *et al*. (Rafnsdóttir et al., 2023), successfully formed spheroids within one week of culture time just like the PHH seeded in FBS-containing medium using the standard protocol. Not only did spheroids form in SFM conditions, but their morphology was also better or equal to the spheroids formed from PHH from the same donor using FBS-supplemented conditions as evaluated by blinded researchers familiar with 3D PHH studies (Supplementary Tables 5 and 6).

Naturally, we observed an inter-donor variability both in the time needed for spheroid formation and in morphology. Yet all these variations were expected and within biological limits, as PHH donor batch variability is a well-known phenomenon. (Ölander et al., 2019, 2020, 2021) This is once again reflected in the morphological score granted by our fellow researchers and our viability measurements.

Further, we continued to track the viability and morphology of 3D PHH from a randomly selected donor (Donor 1) for three weeks. We observed that spheroids formed in both SFM or FBS retained their morphology for all three weeks; however, consistent with previous observations reported in the literature, spheroids formed in FBS exhibited a certain degree of compactization by the second and third week in culture (Figure 2a, Supplementary Figure 2)(Bell et al., 2016; Xing et al., 2024). Additionally, and in line with previous reports, a decrease in overall ATP content/viability was observed in spheroids formed in FBS between the first and second weeks of culture (Bell et al., 2018; Handin et al., 2021; Kanebratt et al., 2021). Surprisingly, we did not observe this phenomenon in 3D PHH formed in SFM, instead they kept steady levels of ATP throughout the observation time.

Interestingly, the CYP data support our observations regarding viability and morphology. Overall, CYP protein expression levels remained more stable in 3D PHH formed in SFM, with the activity of CYP3A4 and CYP2C9 being significantly higher in 3D PHH formed in SFM particularly during the second and the third weeks of culture. A decrease in CYP activity in spheroids formed in FBS could be connected to a decrease in ATP content. Kanebratt *et al*. have previously demonstrated that CYP activity follows along with ATP content, and when CYP metabolites formation rate is normalized to ATP levels – metabolic formation comes across as stable (Kanebratt et al., 2021b).

Considering that only minor differences were seen in morphology and functional performance of 3D PHH formed in FBS-supplemented medium or SFM, we did not expect the proteome to show any major differences between the two conditions. Indeed, global proteomics analysis revealed a high degree of similarity in the proteomics fingerprint of these cultures. In differential expression analysis, throughout the three weeks of culturing, less than three percent of the proteome was significantly different between FBS and SFM conditions. Only one protein, glutathione peroxidase 1 (GPX1), was consistently upregulated in spheroids formed in SFM during all three weeks of culture. GPX1 is an intracellular antioxidant enzyme that reduces hydrogen peroxide and lipid peroxides and prevents oxidative damage. (Lubos et al., 2011) As a selenocysteine-containing enzyme, GPX1 expression is regulated by the levels of selenium and selenocysteine incorporation during protein translation (Lei et al., 2007). Here, we hypothesize, that additional selenium supplementation from the serum-free cocktail (Supplementary Table 2) during the spheroid formation augments the selenium pool in PHH, and therefore increases the expression of GPX1 expression and prevents oxidative stress. At the same time, we observed consistent upregulation of fatty acid-binding protein 5 (FABP5) in 3D PHH formed in FBS, a protein that has previously been demonstrated as a ferroptosis marker in other cell types (Peng et al., 2024). As ferroptosis is a type of programmed cell death that is usually accompanied by an iron-related accumulation of reactive oxygen species (ROS) and decreased antioxidant capacity (Li et al., 2020), these findings together suggest an improved antioxidative machinery function of 3D PHH formed in SFM when compared to spheroids formed in FBS. Of note, this suggestion is based on the observations from one PHH donor, thus these findings warrant further investigation.

Furthermore, we evaluated the expression and function of the ADMET proteins mentioned in the recently adopted (May 2024) IHC M12 guidelines for drug interactions (ICH M12 Guideline on drug interaction studies, 2024). Overall, at given resolution, the proteins listed in the ICH guidelines were stably expressed in both sets of 3D PHH and we observed almost no difference between the two conditions. We noted some significant expression differences in CYP3A5 as well as in UGT1A3, UGT1A9, and UGT2B4. Besides the significant changes seen, an interesting trend was observed with many of the ADMET proteins showing decreasing expression levels over time in 3D PHH formed in FBS whereas the corresponding proteins in SFM-formed 3D PHH did not show the same pattern but instead remained stable over time. This was especially evident for CYP2B6, CYP3A4, CYP3A5 and OATP1B3.

Altogether, our data demonstrates that 3D PHH could be formed in 3R-compliant chemically defined medium, without the addition of FBS, and successfully used for *in vitro* ADMET studies. Morphological and viability evaluations along with global proteomics data, demonstrated that animal serum-free liver *in vitro* spheroid cultures are equal to regular FBS-based *in vitro* models, and sometimes even supersede the FBS-based spheroids, with regard to the viability and functional performance of CYPs. Of note, our long-term studies were performed using only one PHH donor, hence need to be treated with caution until further verification. Yet, we encourage our fellow researchers to consider transitioning *in vitro* cultures to serum-free conditions, which will not only aid in the transparency of research results but also provide a more ethically sound study design. The approach presented in this study and the publications by Rafnsdóttir *et al*. and Weber *et al*. present an opportunity for other researchers for customization of their cell culture medium of choice towards normoglycemic levels of nutrients (Handin et al., 2021) and humanized serum-free conditions (Rafnsdóttir et al., 2023; Weber et al., 2024, 2022). Notable, the serum-free substitute is not only applicable for 3D PHH. Rafnsdóttir *et al*. and Weber *et al*. have successfully used it for culturing a variety of cell lines. Hence, we anticipate that this serum substitute can be used also for other ADMET cell models. Of note, there is an opportunity for further improvements of cell culture medium by substitution of human-derived proteins (e.g. plasma-derived human serum albumin or placenta-derived laminins) to pure recombinant proteins and/or meticulous evaluation of the role of every component in the cell culture medium.

Lastly, we share our global proteomics data according to the Findable, Accessible, Interoperable, and Reusable (FAIR) data principles and offer other researchers to use it for benchmarking of the performance of the serum-free 3D PHH cultures. We hope our study will contribute to more informed and eventually more harmonized protocols for 3D PHH cultures, as well as pave the way to serum-free primary human cultures.

## Data availability

The mass spectrometry proteomics data have been deposited to the ProteomeXchange Consortium via the PRIDE partner repository with the dataset identifier PXD056428 and is publicly available as of the date of publication. This paper does not report an original code. Raw data will be made available on reasonable request from the corresponding author.

## Acknowledgments

We are grateful to Professor Per Artursson for providing financial support, the primary human hepatocytes, the global proteomics infrastructure, and invaluable scientific discussions for this project. Further, we are grateful to Professor Volker Lauschke for valuable discussions. We thank the team of 3D PHH experts who helped with the blinded evaluation of spheroids, namely Alina Meyer, Aurino Kemas, Sabine Willems, Stefania Koutsilieri, and Sonia Youhanna. This work was funded by grants from the Torvald and Britta Gahlin Foundation and the Valborg Jacobsson Foundation awarded to Maria Karlgren and Evgeniya Mickols, grant from the Swedish Fund for Research Without Animal Experiments awarded to Professor Stina Oredsson (F2020-0002), as well as Swedish Research Council grants (2020-01586 and 2017-01951) awarded to Professor Per Artursson.

## References

1 Baker, M., 2016. Reproducibility: Respect your cells! Nature 537, 433–435. 10.1038/537433a

2 Bell, C.C., Dankers, A.C.A., Lauschke, V.M., Sison-Young, R., Jenkins, R., Rowe, C., Goldring, C.E., Park, K., Regan, S.L., Walker, T., Schofield, C., Baze, A., Foster, A.J., Williams, D.P., van de Ven, A.W.M., Jacobs, F., van Houdt, J., Lähteenmäki, T., Snoeys, J., Juhila, S., Richert, L., Ingelman-Sundberg, M., 2018. Comparison of Hepatic 2D Sandwich Cultures and 3D Spheroids for Long-term Toxicity Applications: A Multicenter Study. Toxicol Sci 162, 655–666. 10.1093/toxsci/kfx289

3 Bell, C.C., Hendriks, D.F.G., Moro, S.M.L., Ellis, E., Walsh, J., Renblom, A., Fredriksson Puigvert, L., Dankers, A.C.A., Jacobs, F., Snoeys, J., Sison-Young, R.L., Jenkins, R.E., Nordling, Å., Mkrtchian, S., Park, B.K., Kitteringham, N.R., Goldring, C.E.P., Lauschke, V.M., Ingelman-Sundberg, M., 2016. Characterization of primary human hepatocyte spheroids as a model system for drug-induced liver injury, liver function and disease. Sci Rep 6, 25187. 10.1038/srep25187

4 Bielow, C., Mastrobuoni, G., Kempa, S., 2016. Proteomics Quality Control: Quality Control Software for MaxQuant Results. J Proteome Res 15, 777–787. 10.1021/acs.jproteome.5b00780

5 Cox, J., Mann, M., 2008. MaxQuant enables high peptide identification rates, individualized p.p.b.-range mass accuracies and proteome-wide protein quantification. Nat Biotechnol 26, 1367–1372. 10.1038/nbt.1511

6 Dai, W., Chen, Y., Xiong, W., Li, S., Tan, W.-S., Zhou, Y., 2024. Development of a serum-free medium for myoblasts long-term expansion and 3D culture for cell-based meat. J Food Sci 89, 851–865. 10.1111/1750-3841.16884

7 Didusch, S., Madern, M., Hartl, M., Baccarini, M., 2022. amica: an interactive and user-friendly web-platform for the analysis of proteomics data. BMC Genomics 23, 817. 10.1186/s12864-022-09058-7

8 Eden, E., Lipson, D., Yogev, S., Yakhini, Z., 2007. Discovering Motifs in Ranked Lists of DNA Sequences. PLOS Computational Biology 3, e39. 10.1371/journal.pcbi.0030039

9 Eden, E., Navon, R., Steinfeld, I., Lipson, D., Yakhini, Z., 2009. GOrilla: a tool for discovery and visualization of enriched GO terms in ranked gene lists. BMC Bioinformatics 10, 48. 10.1186/1471-2105-10-48

10 Edwards, A., Roscoe, L., Longmore, C., Bailey, F., Sim, B., Treasure, C., 2018. Adaptation of the human Cell Line Activation Test (h-CLAT) to Animal-Product-Free Conditions. ALTEX 35, 477–488. 10.14573/altex.1710051

11 European Medicines Agency, 2024. ICH M12 on drug interaction studies https://www.ema.europa.eu/en/ich-m12-drug-interaction-studies-scientific-guideline

12 Fiol, C.R., Collignon, M.-L., Welsh, J., Rafiq, Q.A., 2023. Optimizing and developing a scalable, chemically defined, animal component-free lentiviral vector production process in a fixed-bed bioreactor. Mol Ther Methods Clin Dev 30, 221–234. 10.1016/j.omtm.2023.06.011

13 Gheytanchi, E., Naseri, M., Karimi-Busheri, F., Atyabi, F., Mirsharif, E.S., Bozorgmehr, M., Ghods, R., Madjd, Z., 2021. Morphological and molecular characteristics of spheroid formation in HT-29 and Caco-2 colorectal cancer cell lines. Cancer Cell Int 21, 204. 10.1186/s12935-021-01898-9

14 Gstraunthaler, G., Lindl, T., van der Valk, J., 2013. A plea to reduce or replace fetal bovine serum in cell culture media. Cytotechnology 65, 791–793. 10.1007/s10616-013-9633-8

15 Handin, N., Mickols, E., Ölander, M., Rudfeldt, J., Blom, K., Nyberg, F., Senkowski, W., Urdzik, J., Maturi, V., Fryknäs, M., Artursson, P., 2021. Conditions for maintenance of hepatocyte differentiation and function in 3D cultures. iScience 24, 103235. 10.1016/j.isci.2021.103235

16 Hendriks, D.F.G., Vorrink, S.U., Smutny, T., Sim, S.C., Nordling, Å., Ullah, S., Kumondai, M., Jones, B.C., Johansson, I., Andersson, T.B., Lauschke, V.M., Ingelman-Sundberg, M., 2020. Clinically Relevant Cytochrome P450 3A4 Induction Mechanisms and Drug Screening in Three-Dimensional Spheroid Cultures of Primary Human Hepatocytes. Clinical Pharmacology & Therapeutics 108, 844–855. 10.1002/cpt.1860

17 Huber, W., von Heydebreck, A., Sültmann, H., Poustka, A., Vingron, M., 2002. Variance stabilization applied to microarray data calibration and to the quantification of differential expression. Bioinformatics 18 Suppl 1, S96–104. 10.1093/bioinformatics/18.suppl_1.s96

18 Järvinen, E., Hammer, H.S., Pötz, O., Ingelman-Sundberg, M., Stage, T.B., 2023. 3D Spheroid Primary Human Hepatocytes for Prediction of Cytochrome P450 and Drug Transporter Induction. Clin Pharmacol Ther 113, 1284–1294. 10.1002/cpt.2887

19 Kanebratt, K.P., Janefeldt, A., Vilén, L., Vildhede, A., Samuelsson, K., Milton, L., Björkbom, A., Persson, M., Leandersson, C., Andersson, T.B., Hilgendorf, C., 2021a. Primary Human Hepatocyte Spheroid Model as a 3D In Vitro Platform for Metabolism Studies. JPharmSci 110, 422–431. 10.1016/j.xphs.2020.10.043

20 Kanebratt, K.P., Janefeldt, A., Vilén, L., Vildhede, A., Samuelsson, K., Milton, L., Björkbom, A., Persson, M., Leandersson, C., Andersson, T.B., Hilgendorf, C., 2021b. Primary Human Hepatocyte Spheroid Model as a 3D In Vitro Platform for Metabolism Studies. JPharmSci 110, 422–431. 10.1016/j.xphs.2020.10.043

21 Katayama, T., Takechi, M., Murata, Y., Chigi, Y., Yamaguchi, S., Okamura, D., 2024. Development of a chemically disclosed serum-free medium for mouse pluripotent stem cells. Front Bioeng Biotechnol 12, 1390386. 10.3389/fbioe.2024.1390386

22 Koutsilieri, S., Mickols, E., Végvári, Á., Lauschke, V.M., 2024. Proteomic workflows for deep phenotypic profiling of 3D organotypic liver models. Biotechnology Journal 19, 2300684. 10.1002/biot.202300684

23 Kozyra, M., Johansson, I., Nordling, Å., Ullah, S., Lauschke, V.M., Ingelman-Sundberg, M., 2018. Human hepatic 3D spheroids as a model for steatosis and insulin resistance. Sci Rep 8. 10.1038/s41598-018-32722-6

24 Lecluyse, E.L., Alexandre, E., 2010. Isolation and culture of primary hepatocytes from resected human liver tissue. Methods Mol Biol 640, 57–82. 10.1007/978-1-60761-688-7_3

25 Lei, X.G., Cheng, W.-H., McClung, J.P., 2007. Metabolic regulation and function of glutathione peroxidase-1. Annu Rev Nutr 27, 41–61.10.1146/annurev.nutr.27.061406.093716

26 Leung, C.M., de Haan, P., Ronaldson-Bouchard, K et al. A guide to the organ-on-a-chip. Nat Rev Methods Primers 2, 33 (2022). 10.1038/s43586-022-00118-6

27 Li, J., Cao, F., Yin, H., Huang, Z., Lin, Z., Mao, N., Sun, B., Wang, G., 2020. Ferroptosis: past, present and future. Cell Death Dis 11, 1–13. 10.1038/s41419-020-2298-2

28 Liu, S., Yang, W., Li, Y., Sun, C., 2023. Fetal bovine serum, an important factor affecting the reproducibility of cell experiments. Sci Rep 13, 1942. 10.1038/s41598-023-29060-7

29 Lubos, E., Loscalzo, J., Handy, D.E., 2011. Glutathione Peroxidase-1 in Health and Disease: From Molecular Mechanisms to Therapeutic Opportunities. Antioxid Redox Signal 15, 1957–1997. 10.1089/ars.2010.3586

30 Mickols, E., Meyer, A., Handin, N., Stüwe, M., Eriksson, J., Rudfeldt, J., Blom, K., Fryknäs, M., Sellin, M.E., Lauschke, V.M., Karlgren, M., Artursson, P., 2024. OCT1 (SLC22A1) transporter kinetics and regulation in primary human hepatocyte 3D spheroids. Sci Rep 14, 17334. 10.1038/s41598-024-67192-6

31 Ölander, M., Wegler, C., Flörkemeier, I., Treyer, A., Handin, N., Pedersen, J.M., Vildhede, A., Mateus, A., LeCluyse, E.L., Urdzik, J., Artursson, P., 2021. Hepatocyte size fractionation allows dissection of human liver zonation. J Cell Physiol 236, 5885–5894. 10.1002/jcp.30273

32 Ölander, M., Wiśniewski, J.R., Artursson, P., 2020. Cell-type-resolved proteomic analysis of the human liver. Liver Int 40, 1770–1780. 10.1111/liv.14452

33 Ölander, M., Wiśniewski, J.R., Flörkemeier, I., Handin, N., Urdzik, J., Artursson, P., 2019. A simple approach for restoration of differentiation and function in cryopreserved human hepatocytes. Arch Toxicol 93, 819–829. 10.1007/s00204-018-2375-9

34 Oliva-Vilarnau, N., Vorrink, S.U., Büttner, F.A., Heinrich, T., Sensbach, J., Koscielski, I., Wienke, D., Petersson, C., Perrin, D., Lauschke, V.M., 2023. Comparative analysis of YAP/TEAD inhibitors in 2D and 3D cultures of primary human hepatocytes reveals a novel non-canonical mechanism of CYP induction. Biochemical Pharmacology 215, 115755. 10.1016/j.bcp.2023.115755

35 Oliva-Vilarnau, N., Vorrink, S.U., Ingelman-Sundberg, M., Lauschke, V.M., 2020. A 3D Cell Culture Model Identifies Wnt/β-Catenin Mediated Inhibition of p53 as a Critical Step during Human Hepatocyte Regeneration. Adv Sci (Weinh) 7, 2000248. 10.1002/advs.202000248

36 Peng, H., Xin, S., Pfeiffer, S., Müller, C., Merl-Pham, J., Hauck, S.M., Harter, P.N., Spitzer, D., Devraj, K., Varynskyi, B., Arzberger, T., Momma, S., Schick, J.A., 2024. Fatty acid-binding protein 5 is a functional biomarker and indicator of ferroptosis in cerebral hypoxia. Cell Death Dis 15, 1–12. 10.1038/s41419-024-06681-y

37 Perez-Diaz, N., Hoffman, E., Clements, J., Cruickshank, R., Doherty, A., Ebner, D., Elloway, J., Fu, J., Kelsall, J., Millar, V., Saib, O., Scott, A., Woods, I., Hutter, V., 2023. Longitudinal characterization of TK6 cells sequentially adapted to animal product-free, chemically defined culture medium: considerations for genotoxicity studies. Front Toxicol 5, 1177586. 10.3389/ftox.2023.1177586

38 Perez-Riverol, Y., Bai, J., Bandla, C., García-Seisdedos, D., Hewapathirana, S., Kamatchinathan, S., Kundu, D.J., Prakash, A., Frericks-Zipper, A., Eisenacher, M., Walzer, M., Wang, S., Brazma, A., Vizcaíno, J.A., 2021. The PRIDE database resources in 2022: a hub for mass spectrometry-based proteomics evidences. Nucleic Acids Res 50, D543–D552. 10.1093/nar/gkab1038

39 Perez-Riverol, Y., Csordas, A., Bai, J., Bernal-Llinares, M., Hewapathirana, S., Kundu, D.J., Inuganti, A., Griss, J., Mayer, G., Eisenacher, M., Pérez, E., Uszkoreit, J., Pfeuffer, J., Sachsenberg, T., Yılmaz, Ş., Tiwary, S., Cox, J., Audain, E., Walzer, M., Jarnuczak, A.F., Ternent, T., Brazma, A., Vizcaíno, J.A., 2019. The PRIDE database and related tools and resources in 2019: improving support for quantification data. Nucleic Acids Research 47, D442–D450. 10.1093/nar/gky1106

40 Preiss, L.C., Lauschke, V.M., Georgi, K., Petersson, C., 2022. Multi-Well Array Culture of Primary Human Hepatocyte Spheroids for Clearance Extrapolation of Slowly Metabolized Compounds. AAPS J 24, 41. 10.1208/s12248-022-00689-y

41 Puck, T.T., Cieciura, S.J., Robinson, A., 1958. Genetics of somatic mammalian cells. III. Long-term cultivation of euploid cells from human and animal subjects. J Exp Med 108, 945–956. 10.1084/jem.108.6.945

42 Rafnsdóttir, Ó.B., Kiuru, A., Tebäck, M., Friberg, N., Revstedt, P., Zhu, J., Thomasson, S., Czopek, A., Malakpour-Permlid, A., Weber, T., Oredsson, S., 2023. A new animal product free defined medium for 2D and 3D culturing of normal and cancer cells to study cell proliferation and migration as well as dose response to chemical treatment. Toxicology Reports 10, 509–520. 10.1016/j.toxrep.2023.04.001

43 Ranaldi, G., Consalvo, R., Sambuy, Y., Scarino, M.L., 2003. Permeability characteristics of parental and clonal human intestinal Caco-2 cell lines differentiated in serum-supplemented and serum-free media. Toxicol In Vitro 17, 761–767. 10.1016/s0887-2333(03)00095-x

44 Rifampicin and its derivatives: stability, disposition, and affinity towards pregnane X receptor employing 2D and 3D primary human hepatocytes - PubMed [WWW Document], n.d. URL https://pubmed.ncbi.nlm.nih.gov/39179119/ (accessed 8.29.24).

45 Rodrigues, A.F., Fernandes, P., Laske, T., Castro, R., Alves, P.M., Genzel, Y., Coroadinha, A.S., 2020. Cell Bank Origin of MDCK Parental Cells Shapes Adaptation to Serum-Free Suspension Culture and Canine Adenoviral Vector Production. Int J Mol Sci 21, 6111. 10.3390/ijms21176111

46 Ross, A.M., Walsh, D.R., Cahalane, R.M., Marcar, L., Mulvihill, J.J.E., 2021. The effect of serum starvation on tight junctional proteins and barrier formation in Caco-2 cells. Biochem Biophys Rep 27, 101096. 10.1016/j.bbrep.2021.101096

47 Stephenson, E., Jacquet, L., Miere, C., Wood, V., Kadeva, N., Cornwell, G., Codognotto, S., Dajani, Y., Braude, P., Ilic, D., 2012. Derivation and propagation of human embryonic stem cell lines from frozen embryos in an animal product-free environment. Nat Protoc 7, 1366–1381. 10.1038/nprot.2012.080

48 Turner, J., Pound, P., Owen, C., Hutchinson, I., Hop, M., Chau, D.Y.S., Barrios Silva, L.V., Coleman, M., Dubourg, A., Harries, L.W., Hutter, V., Kenna, J.G., Lauschke, V.M., Neuhaus, W., Roper, C., Watkins, P.B., Welch, J., Alvarez, L.R., Taylor, K., 2023. Incorporating new approach methodologies into regulatory nonclinical pharmaceutical safety assessment. ALTEX 40, 519–533. 10.14573/altex.2212081

49 Tyanova S, Temu T, Sinitcyn P, Carlson A, Hein MY, Geiger T, Mann M, Cox J. The Perseus computational platform for comprehensive analysis of (prote)omics data. Nat Methods. 2016 Sep;13(9):731–40. doi: 10.1038/nmeth.3901. Epub 2016 Jun 27. PMID: 27348712.

50 UniProt Consortium. UniProt: the Universal Protein Knowledgebase in 2025. Nucleic Acids Res. 2025 Jan 6;53(D1):D609–D617. doi: 10.1093/nar/gkae1010. PMID: 39552041; PMCID: PMC11701636.

51 Välikangas, T., Suomi, T., Elo, L.L., 2018. A systematic evaluation of normalization methods in quantitative label-free proteomics. Briefings in Bioinformatics 19, 1–11. 10.1093/bib/bbw095

52 Valk, J. van der, Bieback, K., Buta, C., Cochrane, B., Dirks, W.G., Fu, J., Hickman, J.J., Hohensee, C., Kolar, R., Liebsch, M., Pistollato, F., Schulz, M., Thieme, D., Weber, T., Wiest, J., Winkler, S., Gstraunthaler, G., 2018. Fetal bovine serum (FBS): Past – present – future. ALTEX - Alternatives to animal experimentation 35, 99–118. 10.14573/altex.1705101

53 Vorrink, S.U., Zhou, Y., Ingelman-Sundberg, M., Lauschke, V.M., 2018. Prediction of Drug-Induced Hepatotoxicity Using Long-Term Stable Primary Hepatic 3D Spheroid Cultures in Chemically Defined Conditions. Toxicol Sci 163, 655–665. 10.1093/toxsci/kfy058

54 Wang, L., Hu, D., Xu, J., Hu, J., Wang, Y., 2024. Complex in vitro model: A transformative model in drug development and precision medicine. Clin Transl Sci 17, e13695. 10.1111/cts.13695

55 Weber, T., Bajramovic, J., Oredsson, S., 2024. Preparation of a universally usable, animal product free, defined medium for 2D and 3D culturing of normal and cancer cells. MethodsX 12, 102592. 10.1016/j.mex.2024.102592

56 Weber, T., Wiest, J., Oredsson, S., Bieback, K., 2022. Case Studies Exemplifying the Transition to Animal Component-free Cell Culture. Altern Lab Anim 50, 330–338. 10.1177/02611929221117999

57 Wisniewski, J.R., Hein, M.Y., Cox, J., Mann, M., 2014. A “Proteomic Ruler” for Protein Copy Number and Concentration Estimation without Spike-in Standards. Molecular & Cellular Proteomics 13, 3497–3506. 10.1074/mcp.M113.037309

58 Xing, C., Kemas, A., Mickols, E., Klein, K., Artursson, P., Lauschke, V.M., 2024. The choice of ultra-low attachment plates impacts primary human and primary canine hepatocyte spheroid formation, phenotypes, and function. Biotechnology Journal 19, 2300587. 10.1002/biot.202300587

