## Supplementary Figure for "Animal product-free formation and cultivation of 3D primary hepatocyte spheroids"

**Supplementary Table 1.** Medical and demographic information of PHH donors used in this study. Age groups: middle-age adult (31-45), old-aged adults (46-75), seniors (>75).

| Donor | Sex | Age group | Diagnosis | BMI |
|---|---|---|---|---|
| 1 | M | old-aged adult | Colorectal cancer | 22 |
| 2 | F | middle-age adult | Colorectal cancer | 28 |
| 3 | F | middle-age adult | Neuroendocrine tumor | 23 |
| 4 | M | senior | Colorectal cancer | 29 |
| 5 | F | senior | Colorectal cancer | 20 |

**Supplementary Table 2.** The serum-free medium composition.

| | | William´s E normoglycemic medium | | Serum-free supplement | |
|---|---|---|---|---|---|
| | | **WE P04-29050S4** | **Added in house** | **Rafnsdottir et al.** | **Total** |
| | | mg/L | mg/L | mg/L | mg/L |
| **Inorganic salts** | NaHCO3 | 2240 | | | 2240 |
| | CaCl2 * 2H2O | 264.92 | | | 264.92 |
| | $Fe(NO_3)_3$ * 9H2O | 0.0001 | | | 0.0001 |
| | KCl | 400 | | | 400 |
| | CuSO4 | 0.0001 | | | 0.0001 |
| | MgSO4 | 97.67 | | | 97.67 |
| | MgCl2 * 4H2O | 0.0001 | | | 0.0001 |
| | NaCl | 6800 | | | 6800 |
| | NaH2PO4 | 140 | | | 140 |
| | ZnSO4 * 7H2O | 0.0002 | | | 0.0002 |
| | $Na_2SeO_3$/H2SeO3 | | 0.005 | 0.008 | 0.013 |
| **Other** | D-glucose | | 990 | | 990 |
| | Glutathione | 0.05 | | 0.012 | 0.062 |

| | | | | | |
|---|---|---|---|---|---|
| | Sodim Pyruvate | 25 | | | 25 |
| | Dexamethasone | | 0.392 | | 0.392 |
| | Penicillin | | 1000 U/mL | | 1000 U/mL |
| | Streptomycin | | 1000 U/mL | | 1000 U/mL |
| | All-trans retinoic acid | | | 0.025 | 0.025 |
| | Cholesterol | | | 0.05 | 0.05 |
| | Hypoxanthine Na | | | 1.75 | 1.75 |
| | O-Phosphoryl ethanolamine | | | 5 | 5 |
| | Pyruvate | | | 110 | 110 |
| | Ribose | | | 0.125 | 0.125 |
| | Xanthine | | | 0.085 | 0.085 |
| | Uracil | | | 0.075 | 0.075 |
| **Fatty acids** | Linoleic acid | | | 1 | 1 |
| | Methyllinoleat | 0.03 | | | 0.03 |
| | Lipoic acid | | | 0.05 | 0.05 |
| | L-Alanine | 90 | | | 90 |
| | L-Arginine free base | 50 | | | 50 |
| | L-Asparagine * H2O | 20 | | | 20 |
| | L-Aspartic acid | 30 | | | 30 |
| | L-Cysteine | 40 | | | 40 |
| | L-Cystine | 20 | | | 20 |
| | L-Glutamine | | 292.28 | | 292.28 |
| | L-Glutamic acid | 50 | | | 50 |
| | Glycine | 50 | | | 50 |
| | L-Histidine base | 15 | | | 15 |

| | | | | | |
|---|---|---|---|---|---|
| | L-Isoleucine | 50 | | | 50 |
| | L-Leucine | 75 | | | 75 |
| | L-Lysine * HCl | 87.5 | | | 87.5 |
| | L-Methionine | 15 | | | 15 |
| | L-Phenylalanine | 25 | | | 25 |
| | L-Proline | 30 | | | 30 |
| | L-Serine | 10 | | | 10 |
| | L-Treonine | 40 | | | 40 |
| | L-Tryptophan | 10 | | | 10 |
| | L-Typosine | 35 | | | 35 |
| | L-Valine | 50 | | | 50 |
| **Vitamins** | L-Ascorbic Acid | 2 | | 0.012 | 2012 |
| | D(+)-Biotin | 0.5 | | | 0.5 |
| | Calciferol/Ergocalciferol | 0.1 | | 0.025 | 0.125 |
| | D-calcium pantothenate | 1 | | | 1 |
| | Choline chloride | 1.5 | | 3.5 | 5 |
| | Folic acid | 1 | | 0.33 | 1.33 |
| | myo-Inositol | 2 | | 4.5 | 6.5 |
| | Menadion sodium bisulfate | 0.01 | | | 0.01 |
| | Nicotinamide | 1 | | | 1 |
| | Pyridoxal * HCl | 1 | | | 1 |
| | Riboflavin | 0.1 | | | 0.1 |
| | Thiamine * HCl | 1 | | 0.08 | 1.08 |
| | Alfa-tocopherol phosphate | 0.01 | | 0.003 | 0.013 |
| | Vitamin A acetate | 0.1 | | | 0.1 |
| | 4-Aminobenzoic acid | | | 0.012 | |

| | | | | | |
|---|---|---|---|---|---|
| | Vitamin B12 | 0.2 | | 0.35 | 0.55 |
| **Hormones** | Insulin | | 0.00058 | | 0.00058 |
| | Triiodothyronine | | | 0.0000002 | 0.0000002 |
| | 17-beta Estradiol | | | 0.0000005 | 0.0000005 |
| | Hydrocortisone | | | 0.00025 | 0.00025 |
| **Proteins and Growth Factors** | Transferrin | | 5.5 | 50 | 55.5 |
| | bFGF | | | 0.001 | 0.001 |
| | Collagen | | | 0.1 | 0.1 |
| | EGF | | | 0.01 | 0.01 |
| | Fetuin | | | 0.04 | 0.04 |
| | iGF1 | | | 0.005 | 0.005 |
| | Laminin | | | 0.02 | 0.02 |
| | pDGF | | | 0.002 | 0.002 |
| | Vitronectin | | | 0.1 | 0.1 |
| | Human Serum Albumin | | | 1250 | 1250 |

**Supplementary table 3.** Liquid chromatography gradient events. Mobile phase A1: Water and 0.1% (v/v) formic acid. Mobile phase B1: Acetonitrile and 0.1% (v/v) formic acid.

| Time, min | flow rate, µl/min | A1, % | B1, % |
|---|---|---|---|
| Initial | 0.5 | 98 | 2 |
| 0.2 | 0.5 | 98 | 2 |
| 1.2 | 0.5 | 5 | 95 |
| 1.7 | 0.5 | 5 | 95 |
| 1.8 | 0.5 | 98 | 2 |
| 2 | 0.5 | 98 | 2 |

**Supplementary table 4.** Metabolite-specific MS parameters.

| Compound | Parent ion (m/z) | Product ion (m/z) | Cone voltage (V) | Collision voltage (V) |
|---|---|---|---|---|

| | | | | |
|---|---|---|---|---|
| 1-hydroxy midazolam | 342.1 | 203 | 34 | 28 |
| 1-hydroxy bufuralol | 278 | 158.9 | 26 | 22 |
| hydroxy bupropion | 256 | 139 | 13 | 27 |
| 4-hydroxy diclofenac | 312.1 | 230.1 | 22 | 32 |

**Supplementary table 5**. Results of the blinded scoring* of 3D PHH.

| Medium | **SFM** | **FBS** | **SFM** | **FBS** | **SFM** | **FBS** | **SFM** | **FBS** | **SFM** | **FBS** |
|---|---|---|---|---|---|---|---|---|---|---|
| Donor | **D1** | **D1** | **D2** | **D2** | **D3** | **D3** | **D4** | **D4** | **D5** | **D5** |
| *Number when coded* | *1* | *9* | *3* | *5* | *8* | *7* | *4* | *6* | *10* | *2* |
| Researcher 1 | 3 | 4 | 4 | 3 | 5 | 5 | 5 | 3 | 4 | 4 |
| Researcher 2 | 4.5 | 4 | 4 | 3 | 5 | 4.5 | 4.5 | 3 | 4 | 4 |
| Researcher 3 | 4 | 2 | 4 | 2 | 5 | 3.5 | 4 | 1 | 4.5 | 3 |
| Researcher 4 | 4 | 3 | 4 | 1 | 5 | 4 | 5 | 2 | 2 | 3 |
| Researcher 5 | 4.5 | 1.5 | 3.5 | 1.5 | 4 | 2 | 3.5 | 1 | 2.5 | 2 |
| **Average** | 3.8 | 2.8 | 3.8 | 1.8 | 4.5 | 3.6 | 4.5 | 2.0 | 2.8 | 3.0 |

* Excellent 3D PHH morphology (as compactness, clear rim, and no excessive debris) was scored as 5, whereas a score of 1 was indicative of poor compactization and visible cell death.

*In supplementary table 5 one could note that, whilst direction of the rating between SFM and FBS is preserved between researchers, the overall rating differs largely between individual researchers. Thus, we performed inter-rated reliability rating (results in Supplementary table 6).*

**Supplementary table 6.** Results of interrater agreement with kappa reliability rating.

| **Category** | **Kappa** | **SE of kappa** | **95% confidence interval** | **Weighted Kappa** |
|---|---|---|---|---|
| FBS formed spheroids | -0.211 | 0.058 | -0.325 to -0.096 | -0.397 |
| SFM-formed spheroids | -0.100 | 0.074 | -0.244 to 0.044 | -0.108 |

Kappa < 0: No agreement; Kappa between 0.00 and 0.20: Slight agreement; Kappa between 0.21 and 0.40: Fair agreement; Kappa between 0.41 and 0.60: Moderate agreement; Kappa between 0.61 and 0.80: Substantial agreement; Kappa between 0.81 and 1.00: Almost perfect agreement.

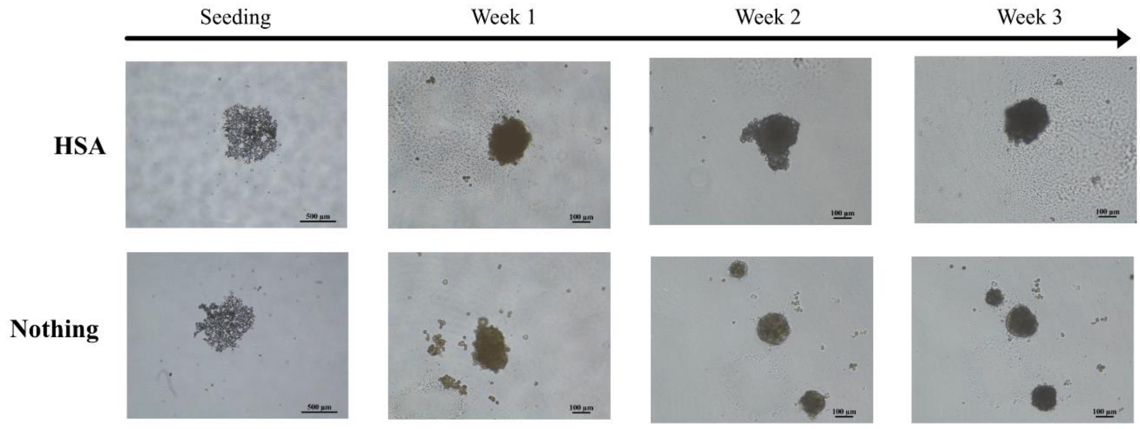


**Supplementary Figure 1.** PHH formation and culturing for 3 weeks in WEng supplemented with only human serum albumin (HSA) or nothing. Note that the majority of the wells did not form spheroids when cultured without FBS or serum-free supplement. The images shown here represent successful spheroid formation under these conditions which only occur for 15-20 % of the wells when cultured with HSA or completely without any supplement.

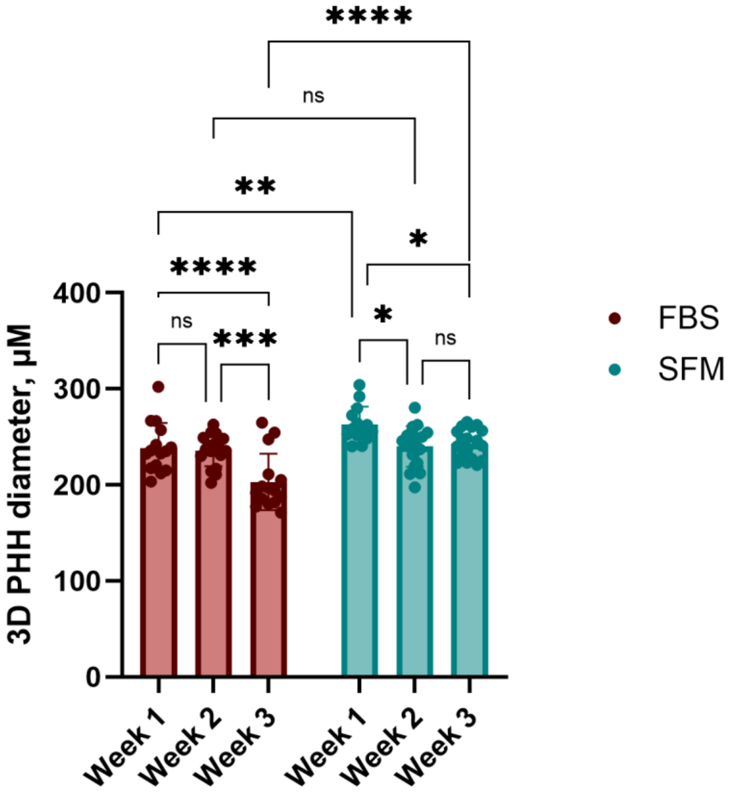


**Supplementary Figure 2.** 3D PHH diameter at every week of culturing. Overall, a certain degree of compactization occurs in both conditions. One could observe a slight decrease in diameter by the third week of culturing in spheroids formed in FBS, while SFM spheroids this trend is less visible. Statistical significance evaluated in two-way ANOVA with Šídák's multiple comparisons test. n = 15 to 19.
